# Distinctive cellular and junctional dynamics independently regulate the rotation and elongation of the internal organ

**DOI:** 10.1101/2023.05.22.541825

**Authors:** Mikiko Inaki, Takamasa Higashi, Satoru Okuda, Kenji Matsuno

## Abstract

Complex structures of organs are formed at high reproducibility, and deformations of epithelia play major roles in these processes. To acquire the intricate morphology, an epithelium simultaneously suffers multiple structural changes. For example, to form the left-right asymmetric structure of *Drosophila* embryonic hindgut, its epithelial tube concurrently rotates and elongates, which are driven by cell sliding and convergent extension, respectively. However, how an epithelium simultaneously accomplishes multiple structural changes remains unclear. To address this issue, we here studied the relevancy between these two mechanisms in the hindgut morphogenesis. Our live imaging analysis revealed that *Myosin1D* and *E*-cad*herin*, or *Par-3* are required only for cell sliding or convergent extension, respectively, while Myosin II is essential for both. Mathematical models showed that these cellular dynamics share a single mechanical system in the same time window. Such specificity and universality of the machineries controlling epithelial dynamics might be a general strategy adopted in complex tissue morphogenesis.

## Introduction

Organ shape is closely related with its functions, so that it needs to be precisely controlled during development. Epithelial morphology often plays major roles to determine the organ shapes. Therefore, the mechanisms of epithelial morphogenesis have been studied extensively, which revealed contributions of, for example, oriented cell division, directional cell migration, polarized cell deformation, and anisotropic cell rearrangement to them ^1–4^. Although the cellular and molecular mechanisms of each individual event in epithelial morphogenesis have been understood well during last decade, in most cases, such mechanisms have been separately studied in simplified systems. However, it is likely that complex morphology of epithelia can be attained only by the cooperative functions of such morphogenetic machineries simultaneously functioning. Considering such prediction, the machineries governing morphogenesis of a tissue can be separated into distinct setups, although they together control the dynamic behaviors of one tissue. However, such complex interplays or independency of these mechanisms have not been understood well.

The *Drosophila* embryonic hindgut is the one of such organs where complex morphogenetic events occur simultaneously. The hindgut is composed of a monolayer epithelium tube and visceral muscles overlaying it ^5^. The embryonic hindgut is firstly formed as bilaterally symmetric structure, whose anterior end curves toward the ventral side of the embryos at stage 12 ^5–7^. Subsequently, two morphogenetic events concurrently proceed at stage 13; the elongation and left-handed rotation of the hindgut epithelial tube, which makes the hindgut a left-right (LR) asymmetric hook-like structure.

The elongation of the hindgut epithelial tube is a typical example of convergent extension ^8^. Similarly, convergent extension is involved in the elongation of the kidney and cochlea in vertebrates and the trachea in *Drosophila* ^9–12^. Convergent extension is also utilized in various epithelial morphogenesis during embryogenesis, such as the germ band extension in *Drosophila* and the dorsal mesoderm extension in *Xenopus* and zebrafish ^13, 14^. During convergent extension, epithelial cells intercalate between each other toward medial direction, which is referred to as cell intercalation, driven by anisotropic cell-junctional remodeling ^2^. Consequently, these tissues become narrower and longer, as observed in the elongation of the *Drosophila* embryonic hindgut ^8^. Genes required for the convergent extension have been revealed by extensive genetic analyses in the *Drosophila* embryonic hindgut, although their functions may not be specific to this process ^15, 16^.

Another event in the morphogenesis of the *Drosophila* embryonic hindgut is its 90-degree counterclockwise rotation as viewed form the posterior. As a result of this rotation, the direction of bending at the anterior tip of the hindgut changes from ventral to right ^7^. Before the rotation begins, the hindgut epithelial cells show chirality manifested itself in LR-asymmetric tilt of their apical surfaces, which is designated as cell chirality ^7, 17, 18^. Dissolution of cell chirality consequently provides a driving force of the hindgut rotation ^7, 19, 20^. This idea was supported by the analyses of various mutations affecting cell chirality and LR asymmetry of the hindgut ^7, 19–22^. For example, in null mutants of *Drosophila Myosin1D* (*Myo1D*), LR asymmetry of various organs, including the rotation direction of the embryonic hindgut, become reversed ^23, 24^. *Myo1D*, also referred to as *Myosin31DF*, encodes a class I myosin ^23, 24^. In *Myo1D* mutants, cell chirality becomes the mirror image of the wild type before the hindgut rotation ^7, 23^. Furthermore, in mutants of *Drosophila E-cadherin* (*E-cad*), encoding an evolutionarily conserved epithelial cell-adhesion protein, the randomization of hindgut-LR asymmetry coincides with the depletion of cell chirality ^7^. As an intermediate step integrating between cell chirality and the hindgut rotation, cell chirality is converted into cell sliding, a novel behavior of epithelial cells, that is, cells change their relative position as sliding in one direction ^20^. Chiral cell sliding is the ultimate step to drive the left-handed rotation of the hindgut ^20^.

Given that convergent extension and chiral cell sliding simultaneously contribute to the morphogenesis of the hindgut epithelium, we found the hindgut excellent system to study potential independency and relevancy between these two morphogenetic machineries. In this study, we conducted live-imaging analyses to detect the effects of various genetic modifications on convergent extension and chiral cell sliding *in vivo*. We also developed an organ culture system of the hindgut with the administration of chemical inhibitors, in which both convergent extension and chiral cell sliding are quantitatively analyzed *ex vivo* at the whole organ and single cell levels. Furthermore, mathematical models of the hindgut morphogenesis were constructed to see respective and integrated contributions of these two machineries to hindgut morphogenesis. These analyses revealed that convergent extension and chiral cell sliding are different events, although they also shear a common component with distinguishable usages in polarized contraction of cell boundaries: anterior-posterior/distal-proximal anisotropy and chirality. We found that the former and the latter independently but concurrently cause convergent extension and chiral cell sliding, respectively. Therefore, our results demonstrate that concurrent actions of distinct morphogenetic machineries are responsible for complex transformation of organ shapes.

## Results

### LR defects associated with *E-cad* and *Myo1D* modifications coincided with cell sliding in the hindgut

During the LR asymmetric development of the embryonic hindgut, the epithelial tube of this organ concurrently rotates and elongates though cell sliding and convergent extension, respectively ^20^. However, it was not clear whether these two epithelial dynamics are operated independently each other or have mutual interdependence. To distinguish between these two possibilities, first, we genetically altered the activities of *E-cad* and *Myo1D* genes that have been shown to play essential roles in the LR-asymmetric development of the hindgut and then examined whether such genetic modifications affect either cell sliding or convergent extension or both.

We previously found the LR asymmetry of the hindgut is mostly randomized in *E-cad* mutants at stage 14 embryos ^7^. In accordance with the previously idea that cell chirality of the hindgut epithelial cells drives the LR-asymmetric rotation of this organ, cell chirality is abolished in these cells of *E-cad* mutants ^7^. Here, to understand the roles of *E-cad* in cell sliding and convergent extension, we performed live imaging of the hindgut in an *E-cad* null mutant, *shotgun^R^*^69^ (*shg^R^*^69^) ^25^. *UAS-Redstinge*r (encoding a nuclear marker) and *UAS-myr-GFP* (encoding a membrane marker) were expressed specifically in the hindgut epithelium by Gal4-UAS system ^26–28^. We used *NP2432* or *byn-gal4* as the hindgut epithelium-specific *Gal4* drivers ^23, 29^. From the dorsal side of the embryo, three-dimensional time-laps images of the hindgut were taken for 120 min from the late stage 12, which is enough for the wild-type hindgut to compete its 90-degree rotation (Fig. 1b1-3, Movie 1) ^20^. However, the hindgut of *shg^R^*^69^ mutant hardly rotated under this condition (Fig. 1a1-3, Movie 2). To visually detect cell sliding, each row of nuclei that lined up along the anterior-posterior axis at 0 min was indicated by different colored circles (Fig. 1d1-3). After 30 and 60 min, the relative positions of nuclei did not change markedly in *shg^R^*^69^ mutant embryos, although the rows of nuclei slanted to the left in wild type, as reported before (Fig. 1d1-3,e1-3) ^20^. Thus, cell sliding was severely diminished in *E-cad* embryos.

**Figure 1:**
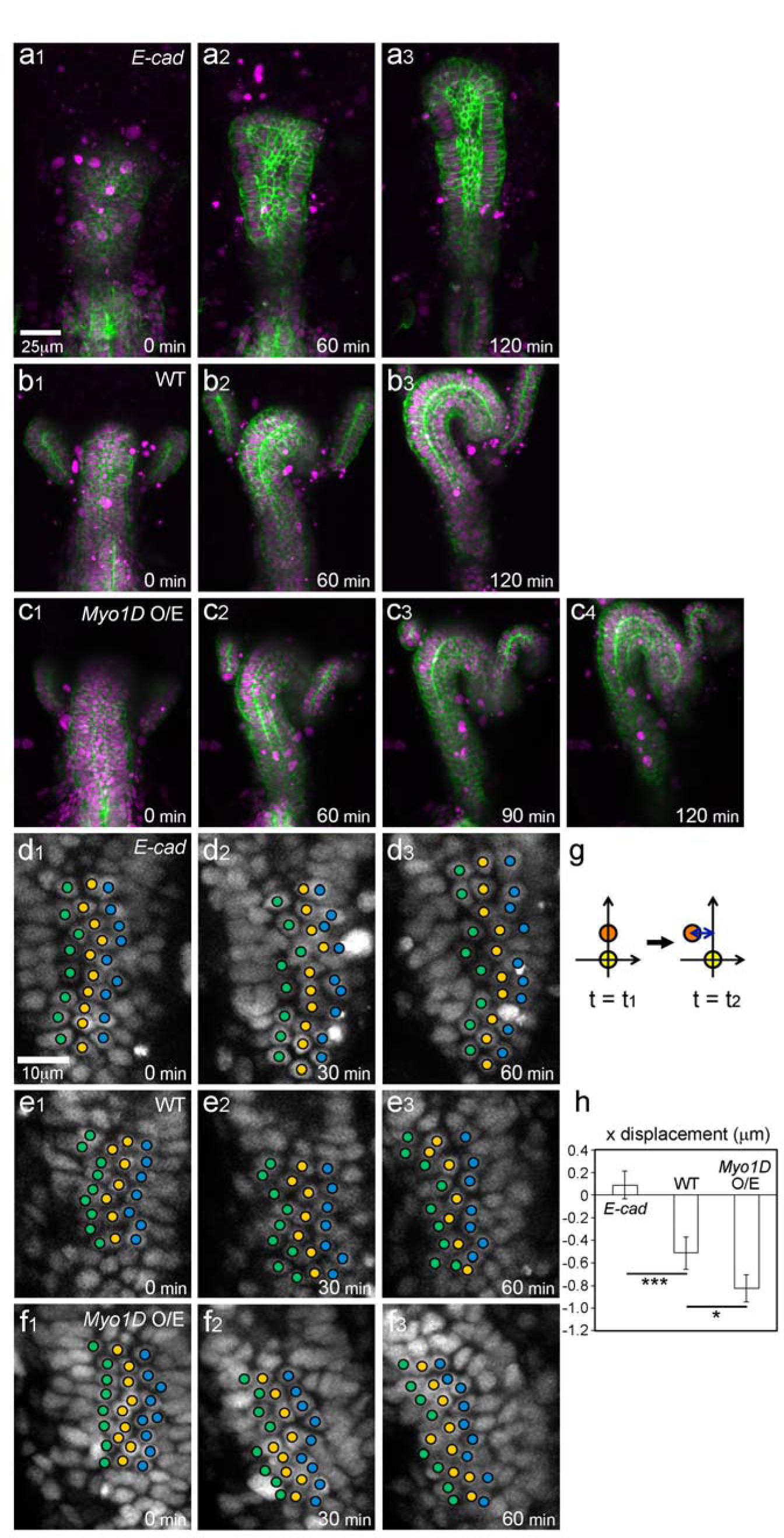
*E-cad* and *Myo1D* perturbation affected the hindgut rotation and cell sliding. (a-c) Still shots from movies of developing *Drosophila* embryonic hindgut visualized by UAS-*myr-GFP* (membrane in green) and UAS-*redstinger* (nuclei in magenta) driven by *byn*-Gal4 during wild-type rotation period (from late stage 12 to stage 13) in *E-cad* mutant (a), wild type (b), and with *Myo1D* overexpression (*Myo1D* O/E) (c). (d-f) Cell movement in the hindgut of *E-cad* mutant (d) and wild type (e) and with *Myo1D* overexpression (f) visualized by UAS-*redstinger*. Three central columns of cells are marked by colored circles. Initial vertical cell columns became tilted leftward in wild type (e3) and with *Myo1D* overexpression (f3) but not in *E-cad* mutant (d3). (g) Schematic of the displacement quantification. The coordinates of two cells located along the anterior-posterior axis at two time points were observed. The subjacent cell was set at (0,0), and the relative displacement of the upper cell in the x direction was measured. (h) X displacement during the hindgut rotation in *E-cad* mutant and wild type and with *Myo1D* overexpression. Cells showed minus directional movement in wild type (n: cell number=164, N: embryo number=7) while cells showed little movement in *E-cad* mutant (n=175, N=7). The movement was significantly enhanced with *Myo1D* overexpression (n=180, N=5). Error bars indicate SEM. ***, *p* < 0.001. *, *p* < 0.05. For all images, anterior is up. Elapsed time from the start of the movie is shown at lower right.

To confirm this observation quantitatively, we measured relative cell displacement in x direction as described previously ^20^. Briefly, we placed a subjacent (posterior in the hindgut) nucleus position at (0, 0) of the coordinates and measured the relative x displacement of the above nucleus against the subjacent nucleus every 30 minutes (Fig. 1g). While wild-type hindgut epithelium had x displacement of minus values showing relative movement leftward as reported previously, the *E-cad* epithelium hardly showed x displacement (Fig. 1h) ^20^. These results demonstrated that cell sliding requires *E-cad*.

*Myo1D* has a dextral activity to define the enantiomorphic states of organ and cell chirality ^7, 23, 24^. Thus, the organ and cell chirality become the mirror image of wild-type counterparts in the absence of *Myo1D* ^7^. In agreement with these findings, the direction of cell sliding is also reversed in *Myo1D* mutant, compared with that of wild type ^20^. In contrast, overexpression of wild-type *Myo1D* induces *de novo* chirality in organs and cells ^23, 30^. Therefore, in this study, we overexpressed a wild-type *Myo1D* in the hindgut epithelium driven by *byn*-Gal4 and analyzed the potential reinforcement of cell sliding. As shown in Fig. 1b, the wild-type hindgut took approximately 120 min to complete 90-degree rotation (Fig. 1b1-3). However, upon the overexpression of *Myo1D*, the hindgut occasionally showed over-rotation (6 out of 13). Under this condition, the hindgut often completed its 90-degree rotation by 90 min and continued to rotate further at least until 120 min (Fig. 1c1-4 and Movie 4).

We found that such acceleration of the hindgut rotation coincided with an increase of cell sliding (Fig. 1f,h). In the *Myo31DF*-overexpression condition, the rows of nuclei at 0 min gradually slanted to the left after 30 and 60 min, which was greater in degree than that of wild type (Fig. 1f1-3). This result was quantitatively supported by the analysis of relative cell displacement in which the x displacement showed significantly larger negative number as compared with that of the wild type (Fig. 1h). These results suggest a positive correlation between the degree of cell sliding and the expression level of *Myo31DF*. Taken the results of *E-cad* and *Myo1D* analyses together, we established conditions to reduce and facilitate the cell sliding, which can be used to test whether cell sliding is coupled with convergent extension.

### *E-cad* and *Myo1D* did not affect convergent extension

We next analyzed whether cell sliding is concomitant with convergent extension in the hindgut epithelium. To quantitatively analyze the degrees of the hindgut rotation and extension simultaneously in a live embryo, we here developed a novel procedure. The hindgut has hook-like structure, and the peak of elbowlJshaped bend in this structure is defined to evaluate the rotation (Fig. 2a). The root of the hook-like structure is largely straight, so that the direction of hindgut lumen is parallel to its rotation axis, which is defined as “pivot line” (Fig. S1). Thus, as the hindgut rotates, the distance between the peak of elbowlJshaped bend and the pivot line increases (Fig. S1). The distance between the pivot line and the center of the hindgut lumen at the peak of elbowlJshaped bend is measured at 0 and 60 min (Fig. 2a). We referred to the change of these values from 0 to 60 min time points as “rotational movement index”. At the same time, to quantify the elongation of the hindgut, a line passing the most anterior peak of the hook-like structure and perpendicular to the pivot line was drown, and the distance between the line and the posterior end of the curved part in the hook-like structure is measured at 0 and 60 min (Fig. 2c, S1). The ratio of the values at 60 min to that of 0 min are designated as “elongation rate index.”

**Figure 2:**
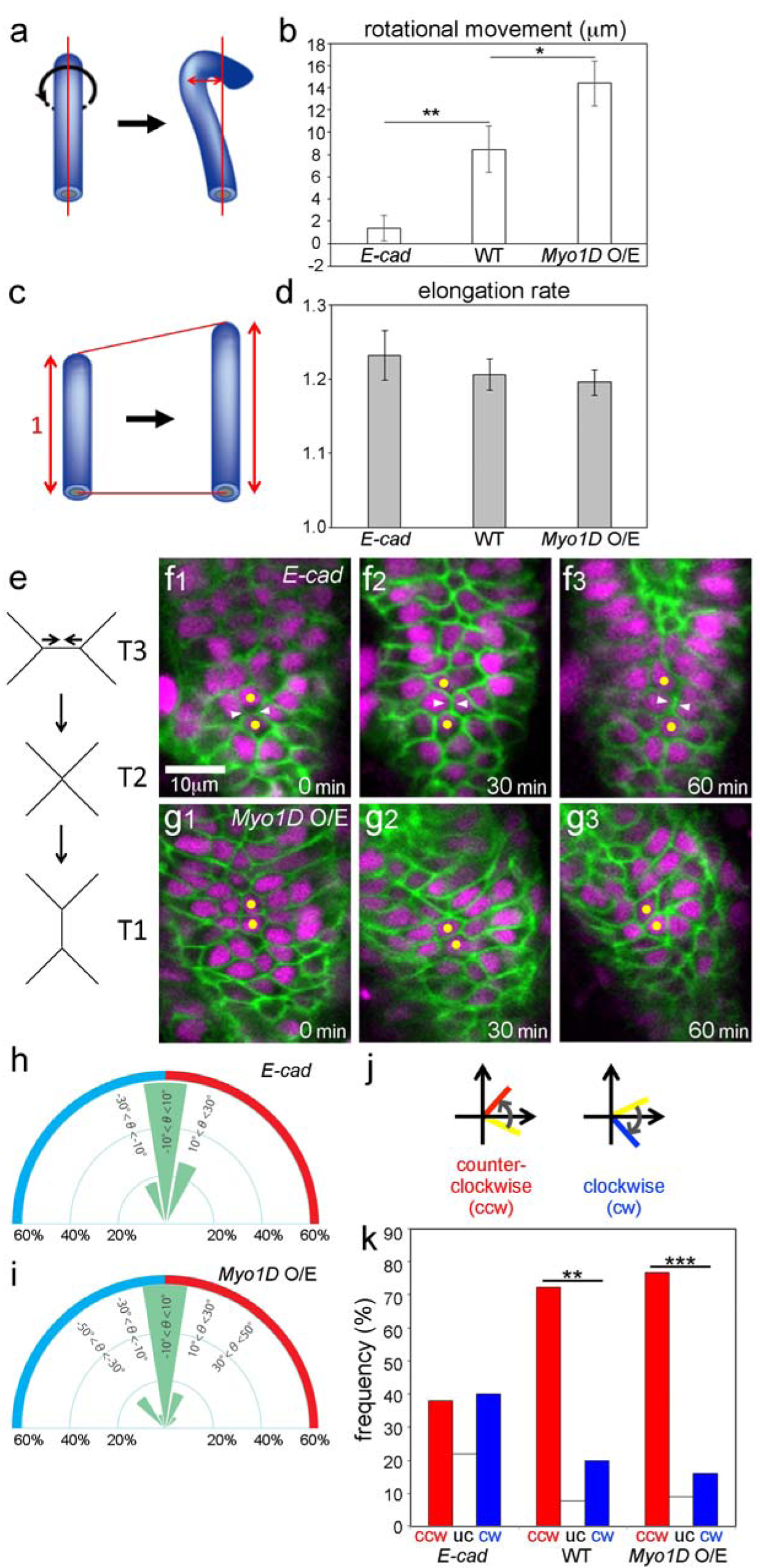
*E-cad* and *Myo1D* perturbation did not affect the hindgut elongation and cell intercalation. (a-d) Quantification of the hindgut deformation. (a,c) Schematic of the rotation (a) and elongation (c) quantification. We measured x displacement of the hindgut peak position by the rotation as rotational movement index and elongation rate with the initial length as 1 during first one hour of the wild-type rotation period. (b,d) Rotational movement (b) and elongation rate (d) indexes in wild type and *E-cad* mutant, and with *Myo1D* overexpression (9≦N≦14). (e) Schematic of T3 to T1 transition. (f,g) Still shots from movies of cell nuclei (magenta) and boundaries (green) in *E-cad*-mutant (f) and *Myo1D*-overexpressed (g) hindgut epithelial cells visualized as in Fig. 1a. Yellow circles indicate cells next to each other in the first frame of the movie. Cells maintained their junction (white arrowheads) during sliding (f). Cells also shows intercalations accompanied by junctional remodeling (f). (h,i) Frequencies of angles of cell boundaries diminished during the junctional remodeling in the hindgut of *E-cad* mutant (h) (n=20, N=4) and with *Myo1D* overexpression (i) (n=12, N=5). (j,k) Angle change of the cell boundaries during the wild-type rotation period. (j) Schemes for boundary tilting. Initial cell boundaries are shown in yellow. Boundaries tilting in a counter-clockwise direction (ccw) are shown in red, while those in a clockwise direction (cw) are in blue. (k) Frequency of boundaries tilting in ccw (red) and cw (blue) and unchanged (uc, white) in the hindgut of wild type and *E-cad* mutant and with *Myo1D* overexpression (50≦n≦65, 5≦N≦7). Error bars indicate SEM. ***, *p* < 0.001. **, *p* < 0.01. *, *p* < 0.05. For all images, anterior is up. Elapsed time from the start of the movie is shown at lower right.

Using this procedure, the rotation and elongation of the hindgut were compared among wild-type and *E-cad* mutant embryos and embryos overexpressing *Myo1D* in the hindgut epithelium. In homozygote of *shg^R^*^69^, the rotational movement index was almost negligible, and the reduction of the value was significant as compared with that of wild type, which coincides with the elimination of cell sliding under this condition as described above (Fig. 2b). However, in these embryos, the elongation rate index was largely comparable to that of wild type (Fig. 2d). Therefore, a correlation between the rotation and elongation of the hindgut was not observed. We also analyzed these values in the hindgut overexpressing the *Myo1D*. The rotation movement index significantly increased under this condition compared with that of wild type, which coincides with the increase of cell sliding as described above (Fig. 2b). In contrast, the elongation rate index did not show marked difference between the wild-type and *Myo1D*-overexpressing embryos, demonstrating the discordance between these two events (Fig.2d). These results suggest that *E-cad* and *Myo1D* affect cell sliding independently of convergent extension.

### *E-cad* and *Myo1D* affect cell sliding independently of cell intercalation

Convergent extension is driven by cell intercalation, observed as a type of anisotropic junctional remodeling, previously designated as T3 to T1 transition (Fig. 2e) ^2^. This transition encourages these epithelia cells to intercalate each other, which consequently leads to the elongation of the hindgut epithelia tube ^8^. Considering above results, we speculated that cell intercalation is not affected neither in *E-cad* mutant embryos or embryos overexpressing *Myo1D*. To test this prediction, we examined the frequencies of the T3 to T1 transition in the hindgut epithelium of these embryos. In wild type, we detected the T3 to T1 transition in 9.3% of cell boundaries in 0-60 min interval, as reported before (Table 1) ^20^. Therefore, most of cell junctions are stably maintained during cell sliding in wild type ^20^. In *shg^R^*^69^ homozygote, in which cell sliding was largely abolished, the T3 to T1 transition was still observed with frequency of 12.7% in the same interval, supporting the independent regulation of these two cellular dynamics (Table1, Fig.2f). Furthermore, in the embryos overexpressing *Myo1D* in their hindgut, the T3 to T1 transition was observed at 7.6% frequency in the same interval, which was largely comparable to that of the wild type (Table1, Fig.2g). These results suggested that the deceleration and acceleration of cell sliding did not affect the frequency of cell intercalation.

Given that the T3 to T1 transition may contribute exclusively to the elongation of the hindgut, we predicted that this anisotropic junctional remodeling should not have LR bias. Previously, we analyzed the angle (θ) between the diminishing cell boundary and the vertical line to the pivot line in wild type and revealed that cell boundaries with angle θ less than 10 degree are selectively diminished, although the angle θ of diminishing cell boundaries did not show detectable LR bias in wild type ^20^. In this study, we analyzed the angle θ of diminishing cell boundaries in *E-cad* mutant embryos and embryos overexpressing *Myo1D*. Despite the significant difference in cell sliding activity in these embryos (Fig. 1h), we did not observe a marked difference in the frequencies of diminishing cell boundaries with every 20-degree angle bin of θ (Fig. 2h,i). Furthermore, we did not detect an LR bias in these frequencies in these two conditions (Fig. 2h,i). These results further suggest that cell sliding is not coupled with cell intercalation.

Although we did not observe marked changes in cell-junctional remodeling upon the modulation of cell sliding, we revealed that a chiral dynamics of cell junctions, detected as the rotation of cell boundary, was coupled with cell sliding but not the junctional remodeling. As describe above, we majored the angle θ of cell boundaries at 0 and 60 min, and the rotational angle of each cell boundary from 0 to 60 min was calculated ^20^. Then, the frequency of cell boundaries (%) that showed counterclockwise or clockwise rotation was obtained (Fig.2j,k) ^20^. We found that the LR bias of the cell-boundary rotation largely disappeared in *shg^R^*^69^ homozygote, compared with that of the wild type (Fig.2k). In contrast, the LR bias of the cell-boundary rotation was augmented in embryos overexpressing *Myo1D*, compared with that of the wild type (Fig.2k). Therefore, cell sliding correlates with the cell-boundary rotation, but not cell-junctional remodeling, under these two genetic conditions. These results further suggest that the cell sliding and cell intercalation are distinct epithelial dynamics, and *E-cad* and *Myo1D* preferentially contribute to the former.

### Bazooka/Par-3 affects convergent extension but not cell sliding

We next analyzed whether genes affecting convergent extension also have roles in cell sliding. Lengyel *et al.* identified various mutants affecting the convergent extension of the *Drosophila* hindgut ^15, 16^. However, since those mutants, including *bowl*, *lines*, and *drumstick*, showed severe defects in the hindgut development, it was difficult to properly evaluate the rotation and elongation of the hindgut using our procedure (Fig. S2a-c). More recently, it also shown that a mutant of *Par-3*/*bazooka* (*baz*), encoding *Drosophila* Par-3, showed defects in convergent extension during the germband elongation ^31^. We here found that embryos homozygous for a classical allele of *baz*, *baz*^4^, a presumptive null mutant, showed elongation defects in the hindgut (Fig.3a) ^32, 33^. About 25% of these embryos had the hindgut with obscure morphology (designated as deformation), which hindered us analyzing its LR asymmetry (Fig. S2d). However, the rest of embryos (about 75%) had the hindgut with hook-like structure, the majority of which showed normal dextral LR-asymmetry, based on its curving direction, even though they occasionally show the delay in the germband retraction as reported before (Fig. S2d,e). We then conducted quantitative analyses of the rotation and elongation of their hindgut. In the hindgut of *baz*^4^ homozygotes, x displacement (cell sliding) and rotational movement index were equivalent to those of wild type, demonstrating the normal rotation of their hindgut (Fig. 3b,d,e). However, their elongation rate index was significantly reduced in *baz*^4^ homozygotes (Fig. 3c,f), which coincided with a severe reduction of T3 to T1 transition (cell intercalation) to 2.0% frequency, as compared with 9.3% in wild type (Table 1). Therefore, Par-3 regulated hindgut elongation though cell intercalation without affecting cell sliding and hindgut rotation. Based on these results, we speculate that cell sling and cell intercalation are concurrently executed *via* two distinct and independent machineries in the hindgut epithelium.

**Figure 3:**
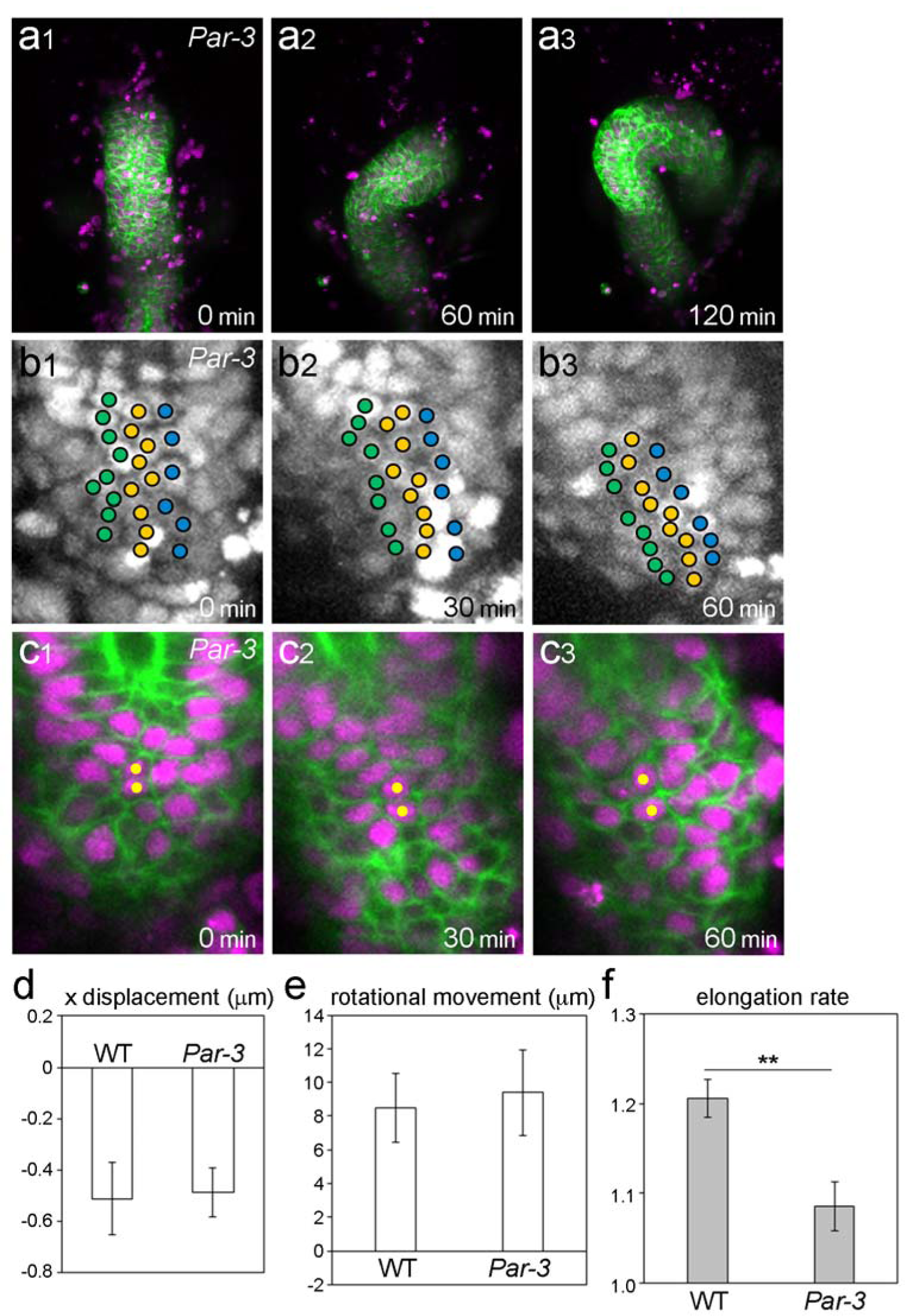
*Par-3* mutants affected the hindgut elongation. (a) Still shots from movies of hindgut rotation in the *Par-3* mutant visualized as in Fig. 1a. (b) Cell movement in the hindgut of *Par-3* mutant visualized as in Fig. 1d. (c) Still shots from a movie of cell nuclei (magenta) and boundaries (green) in *Par-3*-mutant. Yellow circles indicate cells next to each other in the first frame of the movie. (d-f) Quantifications of *Par-3* mutant phenotypes. (d) X displacement of the hindgut epithelial cells in the *Par-3* mutant compared with that in wild type (n=296, N=9). (e,f) Rotational movement (e) and elongation rate (f) indexes in *Par-3* mutant compared with those in wild type (N=7). Error bars indicate SEM. **, *p* < 0.01. For all images, anterior is up. Elapsed time from the start of the movie is shown at lower right.

### Computer model recapitulated that cell sling and convergent extension are independently but concurrently executed though contractile force with distinct polarities

Our analyses showed that cell sliding and convergent extension are independently but concurrently regulated in the hindgut epithelium. To understand underlying mechanisms how these two epithelial dynamics are integrated, we performed numerical simulations using vertex dynamics models ^34–37^. Previously, it was shown that cell sliding and convergent extension rely on two distinct contractions of cell boundaries; contractions with chirality and anterior-posterior/distal-proximal anisotropy are responsible for cell sliding and convergent extension, respectively ^2, 20^. Therefore, we hypothesize that these two contractions may account for the rotation and extension of the hindgut, which occur independently of each other but simultaneously (Fig.4a-c).

**Figure 4:**
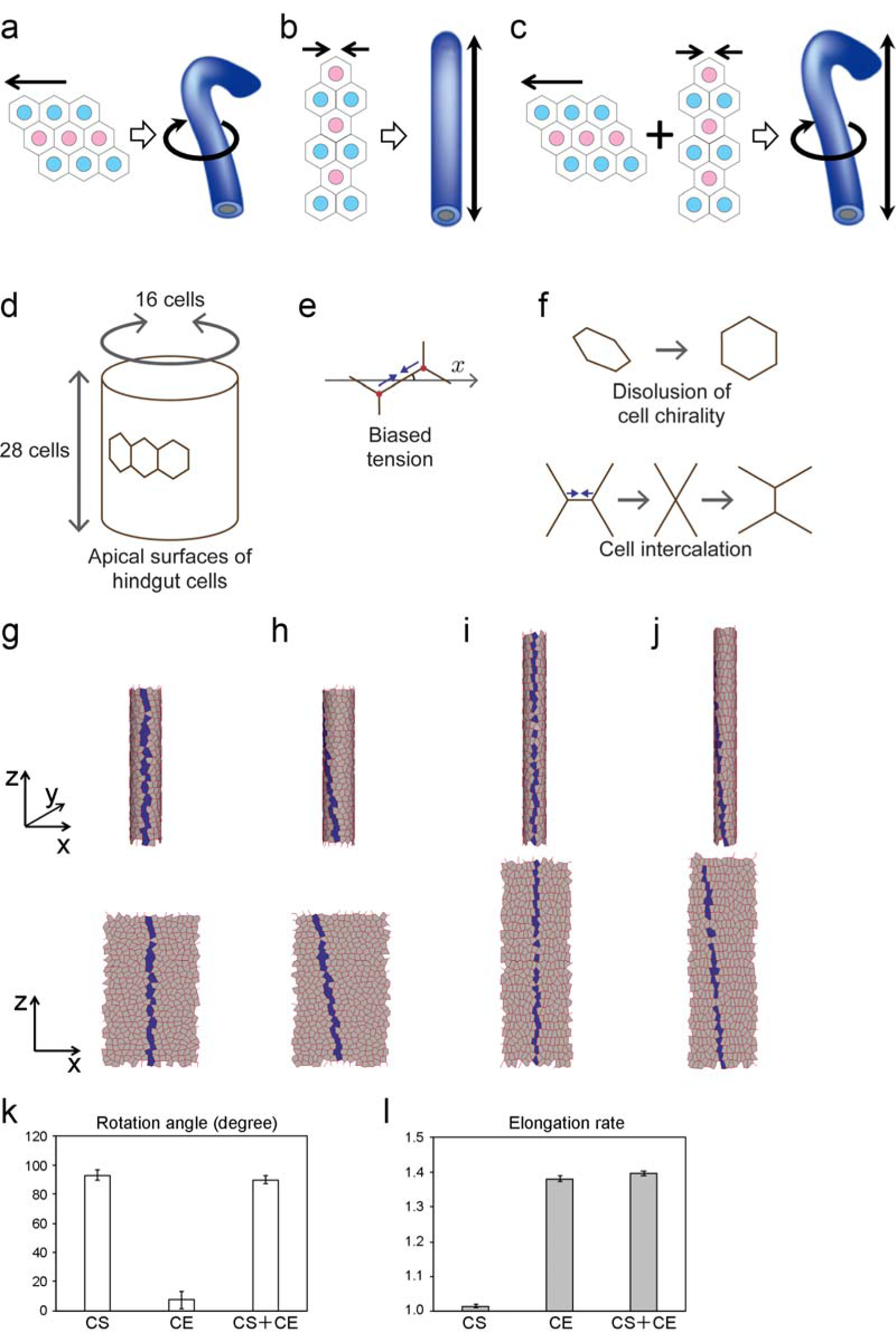
Simulations verified independent and integrated regulation of the hindgut morphogenesis. (a-c) Schemas of cell sliding (a) and cell intercalation (b) regulates the hindgut rotation and elongation, respectively and both cause the hindgut rotation and elongation at the same time (c). (d-e) Schemas of vertex dynamics model. (d) Model tube represents the apical surfaces of the hindgut epithelial cells, which are about 16 cells in circle and 28 cells in length. (e) Biased contraction dependent on edge angle was introduced. (f) For cell sliding, initial cell chirality and subsequent dissolution was introduced (upper). For cell intercalation, horizontally biased edge contraction was introduced, which occasionally result in T1 transition of edges. (g-j) Simulation results. Initial condition with cell chirality (g) and final shapes (h-j) of model tube in 3D (upper) and 2D (lower) resulted from simulation with cell sliding only (h), cell intercalation only (i), and both at the same time (j). (k,l) Quantification of the simulation results. (k) Rotation angle. (l) Elongation rate. SC: cell sliding, CE: convergent extension.

Our previous studies showed that introducing cell chirality by contracting rightward tilted cell edges followed by dissolving this chirality induce cell sliding, which consequently leads to the rotation of hindgut ^7, 20^. In this study, we improved this model by additionally taking axial and radial expansion and contraction of the hindgut into account, which permits the elongation of the hindgut (see Supplementary Information). Briefly, a regular cylinder with 482 regular hexagons (16 in circle and 40 in length, area is 1) was created as a model tube that represents the apical (luminal) surfaces corresponding to the straight region of the hindgut (Fig. 4d). Anterior-posterior/distal-proximal anisotropy in contraction of cell boundaries was introduced as edge tension biased at two oriented angles (Fig. 4e). Cell chirality was introduced as the strain of cell shape under the initial condition, which was generated by the biased edge tension that is maximal at-45 degrees from the *x*-axis. During the simulation, this tension was dissolved, and a horizontal biased edge tension that is maximal at 0 degree was added (Fig. 4f).

To our current model, cell-boundary contractions with chirality or anterior-posterior/distal-proximal anisotropy was separately introduced. We found that introducing chiral tension and its dissolution caused the model tube to rotate about 90 degrees in the counter-clockwise direction (Fig. 4g). In addition, edge tension with anterior-posterior/distal-proximal anisotropy caused the model tube to elongate in the axial direction (Fig. 4h). Importantly, integrating both tensions caused the tube rotation and elongation at the same time, which recapitulates the hindgut morphogenesis *in vivo* (Fig. 4i).

The deformations of model tubes were quantified by averaging the results from independent 5 simulations for each condition. Incorporation of cell chirality, with or without the anterior-posterior/distal-proximal anisotropic tension, induced about 90-degrees of rotation, while the anterior-posterior/distal-proximal anisotropic tension alone caused almost no rotation (Fig. 4j). Instead, the anterior-posterior/distal-proximal anisotropic tension induced about 1.4-fold axial elongation, with or without the incorporation of cell chirality, while cell chirality alone did not cause axial elongation (Fig. 4k). Considering that the wild-type hindgut tube elongated about 1.2-fold for one hour (Fig. 2d) and rotated 90-degree for two hours (Fig. 1b) *in vivo*, our *in silico* models quantitatively agreed with the *in vivo* behaviors of the hindgut tube, further supporting our hypothesis, i.e., contraction forces with distinct polarities caused two independent tissue deformations, which are integrated by occurring in the same developmental time window.

### MyoII contributes to both cell sliding and cell intercalation

Our *in silico* models predicted that contraction forces with distinct polarities, chirality and anterior-posterior/distal-proximal anisotropy, are crucial for the rotation and elongation of the hindgut. Although it is difficult to specifically inhibit only one of these two contraction forces, it is known that MyoII regulates contractile force of cell boundaries in various epithelial dynamics ^2, 4, 38^. Therefore, we evaluated our *in silico* model by inhibiting MyoII and examining its effects on the rotation and elongation of the hindgut. The MyoII heavy chain (MHC) is encoded by *zipper* (*zip*) gene, and we performed live-imaging of the embryonic hindgut homozygous for *zip*^2^, a null allele of *zip* ^39^. However, *zip*^2^ homozygous embryo showed normal hindgut rotation and elongation, probably because maternal supply of *zip* is sufficient for supporting the normal morphogenesis of this organ (Fig. S3a). Thus, to further inhibit *MyoII*, we misexpressed a dominant-negative form of *zip*, *DN-zip* in the hindgut epithelium of *zip*^2^ heterozygous background ^40^. The LR inversion and bilateral phenotypes of their hindgut were observed, suggesting that MHC is required for proper hindgut rotation (Fig. S3b,c). We also examined a null mutant of *spaghetti squash* (*sqh*), which encodes Myosin regulatory right chain (MRLC) ^41^. In embryos homozygous for *sqh^AX^*^3^, a null allele of *sqh* ^42^, the LR-inversion and no laterality phenotypes of the hindgut were observed, suggesting that MRLC is also important for the hindgut rotation (Fig.S3b). However, the percentages of LR defects in these mutant conditions were too low to conduct quantitative analyses at a cell level, probably due to the residual activity of MyoII.

To solve this problem, we here developed an *ex vivo* assay, which was derived from an *ex vivo* organ culture system previously reported ^20^. Briefly, we dissected the most posterior part of embryo, which is mainly composed of the hindgut, before the initiation of its rotation ^20^. In our new system, the cultured hindgut can be treated with chemical inhibitors and its morphological changes are quantitatively analyzed *ex vivo* (Fig. 5). We found that the hindgut rotated and elongated *ex vivo* as observed *in vivo* (Movie 9). Using this *ex vivo* system, we administered an inhibitor against MyoII, Y-27632, which blocks a MyoII activator, RhoK, to the cultured hindgut (Movie 10)^43^. The rotational movement index was obtained *ex vivo* under this condition using the same procedures conduced *in vivo* as described above. We found that the rotational movement index was almost zero in the Y-27632 treated wild-type hindgut *ex vivo*, whereas the mock-treated hindgut (control) showed this index equivalent to that of wild type *in vivo* (Fig. 5g). Therefore, the hindgut rotation was diminished by the suppression of MyoII activity. We then analyzed the effect of Y-27632 treatment on cell sliding. In the mock-treated hindgut cultured *ex vivo*, the rows of nuclei lined up along the anterior-posterior axis at 0 min slanted to the left after 30 and 60 min, as found in wild type *in vivo* (compare Fig. 1e1-3 and Fig. 5b1-3). However, the rows of nuclei in the wild-type hindgut treated with Y-27632 *ex vivo* did not change their direction for 60 min, demonstrating that cell sliding was abolished (Fig. 5c1-3). This result was quantitatively supported by our findings that x displacement was negligible in the wild-type hindgut with Y-27632 treatment *ex vivo*, whereas the mock-treated hindgut *ex vivo* showed x displacement largely equivalent to that of wild type *in vivo* (Fig.5f). These results suggested that an activity of MyoII is essential for cell sliding.

**Figure 5:**
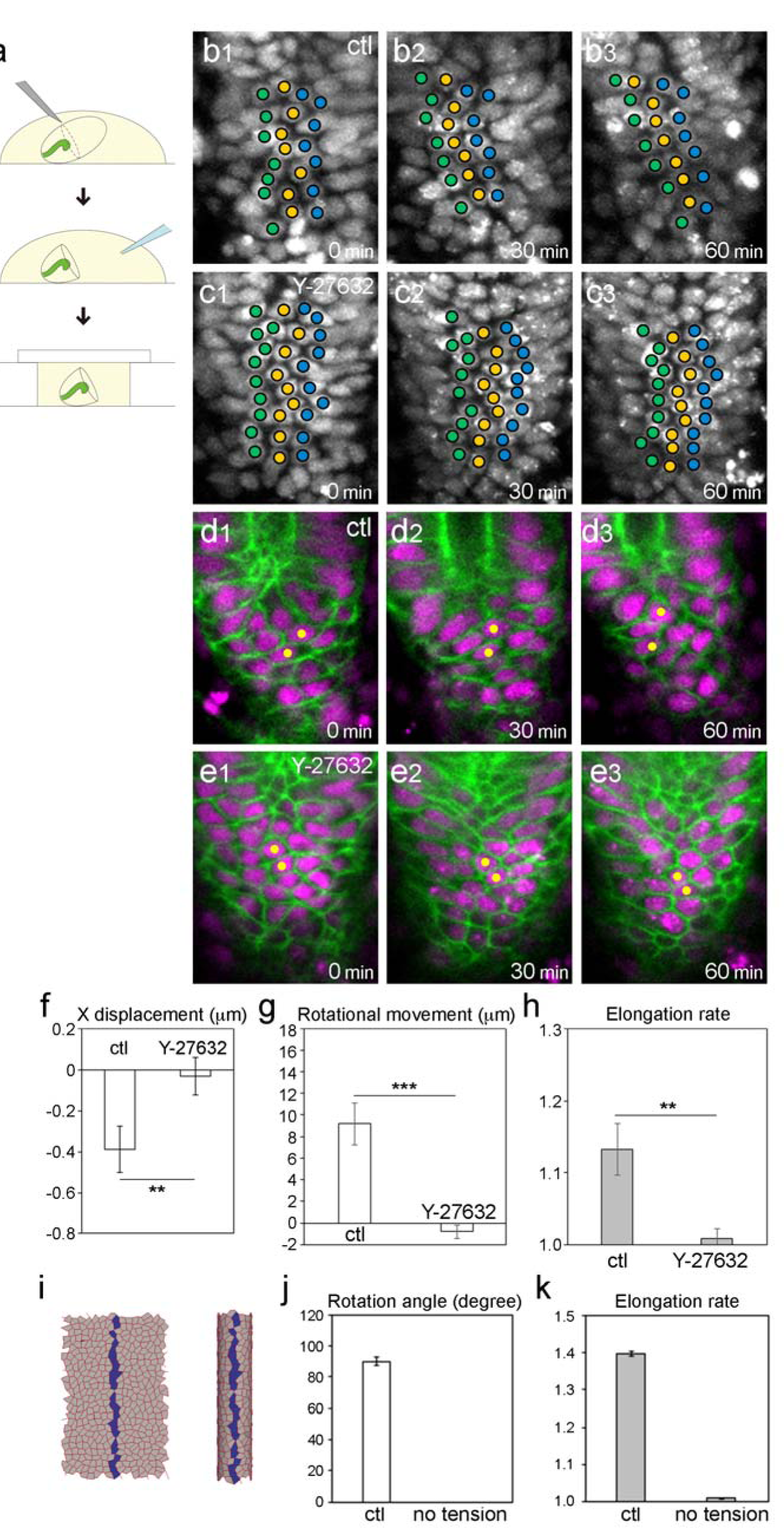
Inhibition of Myosin II activity stopped the rotation and elongation of the hindgut. (a) Schema of *ex vivo* drug treatment experiments. Late-stage12 embryos were dissected by glass pipette needle, treated with a drug or DMSO (control), and encapsulated with a coverslip. (b,c) Cell movement in the *ex vivo* cultured hindgut visualized as in Fig. 1d. Cells slid in DMSO treated control (b) while they did not in Y-27632 treated hindgut (c). (d,e) Still shots from a movie of cell nuclei (magenta) and boundaries (green) in the *ex vivo* cultured hindgut. Cells showed junctional remodeling in DMSO treated control (d) while they did not in Y-27632 treated hindgut (e). (f-h) Quantifications of phenotypes caused by the drug treatment. (f) X displacement of the hindgut epithelial cells in Y-27632 treated compared with that in DMSO treated (155≦n≦232, N=5). (g,h) Rotational movement (g) and elongation rate (h) indexes in Y-27632 treated compared with those in DMSO treated (7≦N≦9). (i-k) Simulation results. No rotation or elongation was observer in the simulation with removal of edge tension after the initial chirality introduction. (i) Final shape of model tube in 3D (right) and 2D (left). (j,k) Quantification of the simulation results. (j) Rotation angle. (k) Elongation rate. Error bars indicate SEM. ***, *p* < 0.001. **, *p* < 0.01. For all images, anterior is up. Elapsed time from the start of the movie is shown at lower right.

Using the same *ex vivo* assay, the requirements of MyoII for the hindgut elongation was also tested. We found that the elongation rate index was close to 1 (no elongation) in the Y-27632 treated wild-type hindgut *ex vivo*, whereas the mock-treated hindgut elongated to a similar extent as that of wild type *in vivo* (Fig. 5h). There results suggested that an activity of MyoII is also essential for the convergent extension of the hindgut. Therefore, we predicted that the T3 to T1 cell-junctional remodeling, which drives convergent extension, may be impaired under this condition. We found that the T3 to T1 transition was severely inhibited by the addition of Y-27632, as compared with that of mock-treated wild type *ex vivo* (Table 2, Fig. 5d,e). Collectively, our results suggested that MyoII-dependent activity, which is presumed to be contraction forces, is required for both cell sliding and convergent extension. This idea agrees with the predictions from our *in silico* model.

To validate this idea more directly in our *in silico* model, we abolished dissolution of cell chirality and anterior-posterior/distal-proximal anisotropic contractions by removing edge tensions from the model, although chirality was introduced before the removal of edge tensions (Fig. 5i). We found that this model gut did not showed rotation nor elongation (Fig. 5i-k), which is reminiscent of the hindgut where the activity of MyoII was inhibited *ex vivo* (Fig. 5c,e,g,h). Taken together, our results suggested that cell sliding and convergent extension are distinct epithelial dynamics induced by chiral and anterior-posterior/distal-proximal anisotropic contractions, which are operated parallelly in the hindgut epithelium at the same developmental time window.

## Discussion

Epithelial morphogenesis often involves multiple types of deformations that concurrently occur. For example, in *Drosophila*, the late stage of gastrulation is coupled with the convergent extension of the germband elongation. Some of pioneer studies revealed that the gastrulation and germband elongation are separable events, and such idea facilities the analyses of specific machineries responsible for each of these events ^2, 14, 38^. However, in general, it is still difficult to separate plural morphogenetic events that take place simultaneously in an epithelium, because we need to find proper objectives that enable us to specifically detect the effects induce by the experimental modulations of each event. To address this, we developed a procedure to analyze a multiplex epithelial dynamics of the *Drosophila* embryonic hindgut, which involves two distinct events simultaneously occur: cell sliding and convergent extension ^8, 20^.

Our analysis revealed that cell sliding in the hindgut epithelial tube depends on *Myo1D* and *E-cad*. In contrast, we failed to detect the requirement of these two genes for the convergent extension, revealed by anisotropic cell-junctional remodeling, although it takes place at the same time as the cell sliding. On the other hand, the convergent extension depends on *Par-3*, which was not required for cell sliding. These results suggested that cell sliding and convergent extension are distinct dynamics of epithelial cells independently operated each other, although they function concurrently in the same epithelium. On the other hand, MyoII is required for both cell sliding and convergent extension of this tissue at the same developmental period. Therefore, although cell sliding and convergent extension are distinct cellular dynamics, they share MyoII as a common key factor, presumably through the control of contraction forces. Our mathematical model revealed that the cell sliding and convergent extension occur at the same time window by using single mechanical system. Although the timing of the anisotropic edge contraction is different: the anisotropic edge contraction for the cell sliding occurs before the rotation while that for the convergent extension occurs after the starting of the rotation, the relaxation process of the cell chirality also depends on edge contractility. Thus, compatibility of these two processes depending on edge contraction was proven by the theoretical model.

Although our results revealed that cell sliding and cell-junctional remodeling are mechanistically separable in the hindgut epithelium, it was previously reported that cell chirality is couple with the chiral distribution of MyoII and LR-asymmetric cell-boundary remodeling during the LR-directional rotation of the male genitalia epithelium ^44^. A computer simulation demonstrated that LR-asymmetric cell-junctional remodeling drives the rotation of doughnut-shaped epithelium ^44^. In contrast, in a vertex model that recapitulates the LR-directional rotation of the hindgut epithelial tube, the dissolution of cell chirality drives the tube rotation, whereas cell-junctional remodeling is dispensable for it ^20^. Therefore, it is determined in organ-specific manners whether cell chirality is couple with cell-junctional remodeling or not, although mechanisms for switching between these two states remain to be understood.

In this study, we developed an organ culture system with the administration of chemical inhibitors to quantitatively analyze the cell sliding, cell-junctional remodeling, and the deformation of the hindgut. This system is particularly useful to study the functions of genes with maternal effect, for example, *zip* and *sqh* ^45^. Maternal contribution of these genes covers the zygotic mutant phenotypes of them. Besides that, complete elimination of gene functions in early embryos, which can be achieved by introducing the maternal and zygotic mutation, often results in the developmental arrest and severe defects at the early stage of embryogenesis. Thus, it makes also difficult to analyze the embryonic organs at a certain time window of development. As an alternative method to suppress the gene functions, RNA interference is generally effective technique in *Drosophila* ^46^. However, tissue-specific RNA interference is often inefficient in embryonic tissues including the hindgut. Thus, it has been difficult to observed the cellular dynamics in the hindgut epithelium in which the functions of a gene are tissue-specifically abolished. In contrast, our organ culture system enabled to temporarily control the time window for suppressing the gene activity by adjusting the timing to add inhibitors. Therefore, on the condition that an inhibitor is available, our organ culture system should provide opportunities to uncover the hidden roles of genes in cell sliding and convergent extension.

## Materials and Methods

### Fly strains

Fly strains used were: *shg^R^*^69 25^, *zip*^2^ (BL8739)^39^, *sqh^AX^*^3^ (BL25712)^42^, *baz*^4^ (BL3295)^32, 33^, *byn*-gal4 ^29^, NP2432-gal4 (Kyoto 104201), UAS-*myrGFP-p10* (JFRC29)^28^, UAS-*redstinger* (BL8547), UAS-*Myo31DF* ^23^, UAS-*DN-zip* ^40^. We raise them at 18 or 25 degrees. Genotype used for live-imaging of *shg* and *baz* mutants and Myo31DF over-expression were *shg^R^*^69^/*shg^R^*^69^, NP2432; UAS-*redstinger*/UAS-*myrGFP-p10*, *baz*^4^/*baz*^4^; UAS-*redstinger*/*byn*-gal4, UAS-*myrGFP-p10* and UAS-*Myo31DF*/+; UAS-*redstinger*/*byn*-gal4, UAS-*myrGFP-p10*, respectively. For selection of *shg^R^*^69^ and *zip*^2^ mutants, we used Cyo-sChFP (BL35523) and Cyo-GFP (BL5195). For selection of *baz*^4^ mutant, FM7c-GFP (BL5193) was used.

### Live imaging and quantification

The live imaging of the hindgut was performed as described previously ^20^. Late stage12 embryos of appropriate genotype were mounted dorsal side up in oxygen-permeable Halocarbon oil 27 (Sigma), using 0.17-0.25 μm-thick coverslips as spacers. We imaged embryos every 5 min for 2 hr with a scanning laser confocal microscope, LSM700 (Zeiss) at 22-25 degrees. For quantification of the cell movement, we tracked cells by their nuclear position manually and measured their x,y coodinates every 30 min until 60 min after rotation began using ImageJ software. We analyzed root part of the hindgut, 20-40% from the bottom, and central three columns of the cells to avoid structural effect on cell movement results ^20^. We set the subjacent cell position at (0,0) and quantified the relative displacement of the upper cell in the x direction (Fig. 1B). For quantification of organ deformation, we measured gut length in anterior-posterior direction and position of hook peak of the hindgut from midline of the embryo at 0 and 60 min after rotation began (Fig. S2). For elongation rate index, the gut length of 60 min was divided by 0min gut length. For rotational movement index, distance between hook peak and midline in x direction at 0min was subtracted from 60 min distance. We used Student’s t-test for statistics.

### *ex vivo* culture and drug treatment

We dissected and culture dechorionated embryos in M3 medium with 10% serum and 1% trehalose on double sticky tape on a slide glass. For MyosinII inhibition, after dissection, Y-27632 (Sigma) dissolved in water was added into culture medium in final concentration of 100μM. We put a coverslip on top of the dissected embryos using 0.17-0.25 μm-thick coverslips as spacers. We took and analyzed images as described above.

### Videos

To track the movement of nuclei, we adjusted the depth of the z stack at each time point to make clear time-lapse movies, because the hindgut cells change their z position over time.

Vertex dynamics model

See Supplementary Information.

---

**Figure S1:**
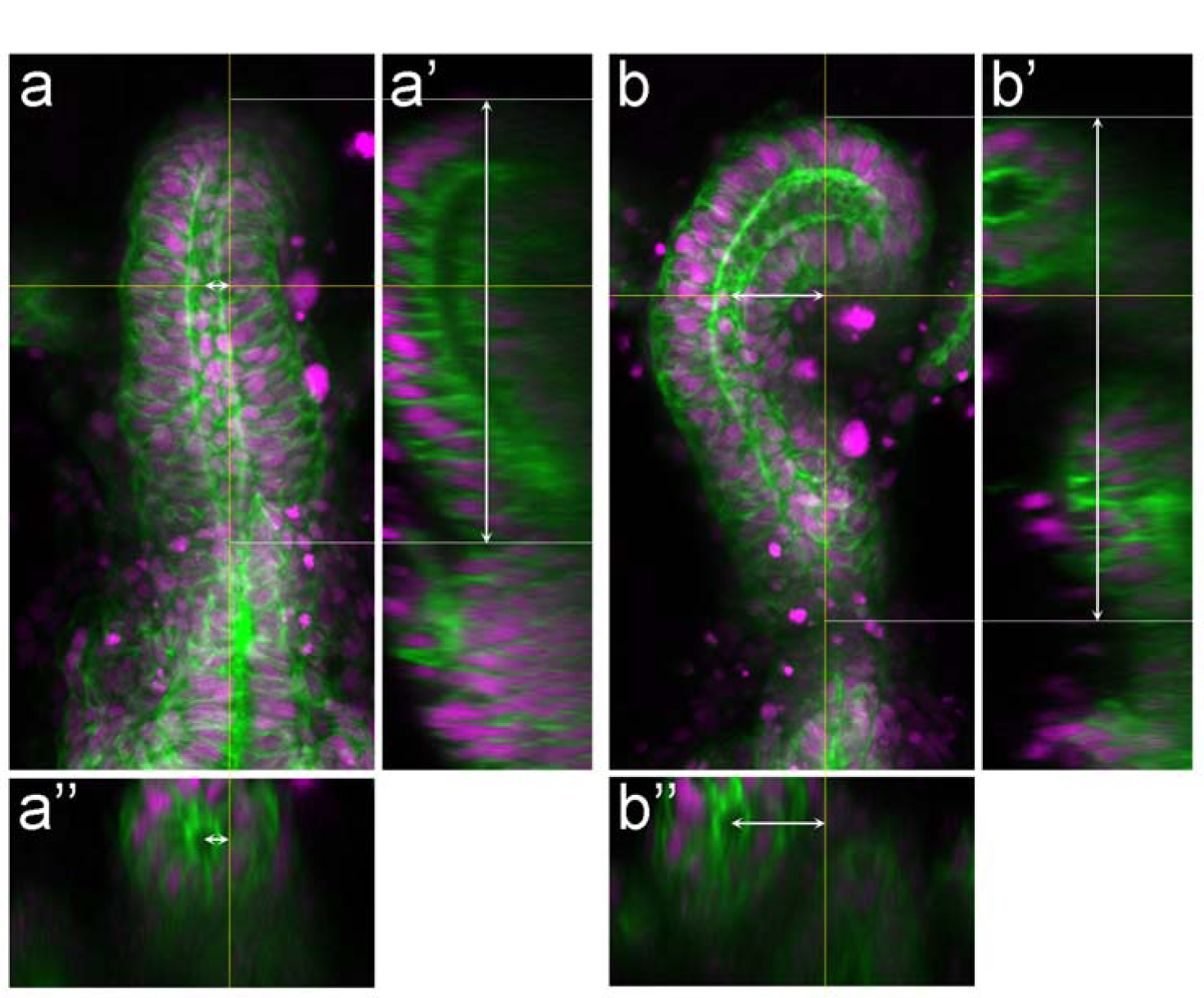
Measurement of the rotational movement and elongation ration indexes. x-y (a,b), y-z (a’,b’) and z-x (a’’,b’’) views of the hindgut at 0 min (a) and 60 min (b) after the movie starts. Pivot lines were drawn from the middle of the lumen of the hindgut bottom part (vertical white lines). The distance the peak of elbow-shaped bend and the pivot line was measured (horizontal white lines with arrowheads), and difference between 0-min and 60-min length was defined as rotational movement index. The heights of the hindgut were also measured (vertical white lines with arrowheads). The elongation ratio of 60-min height with 0-min height as1was defined as elongation ratio index.

**Figure S2:**
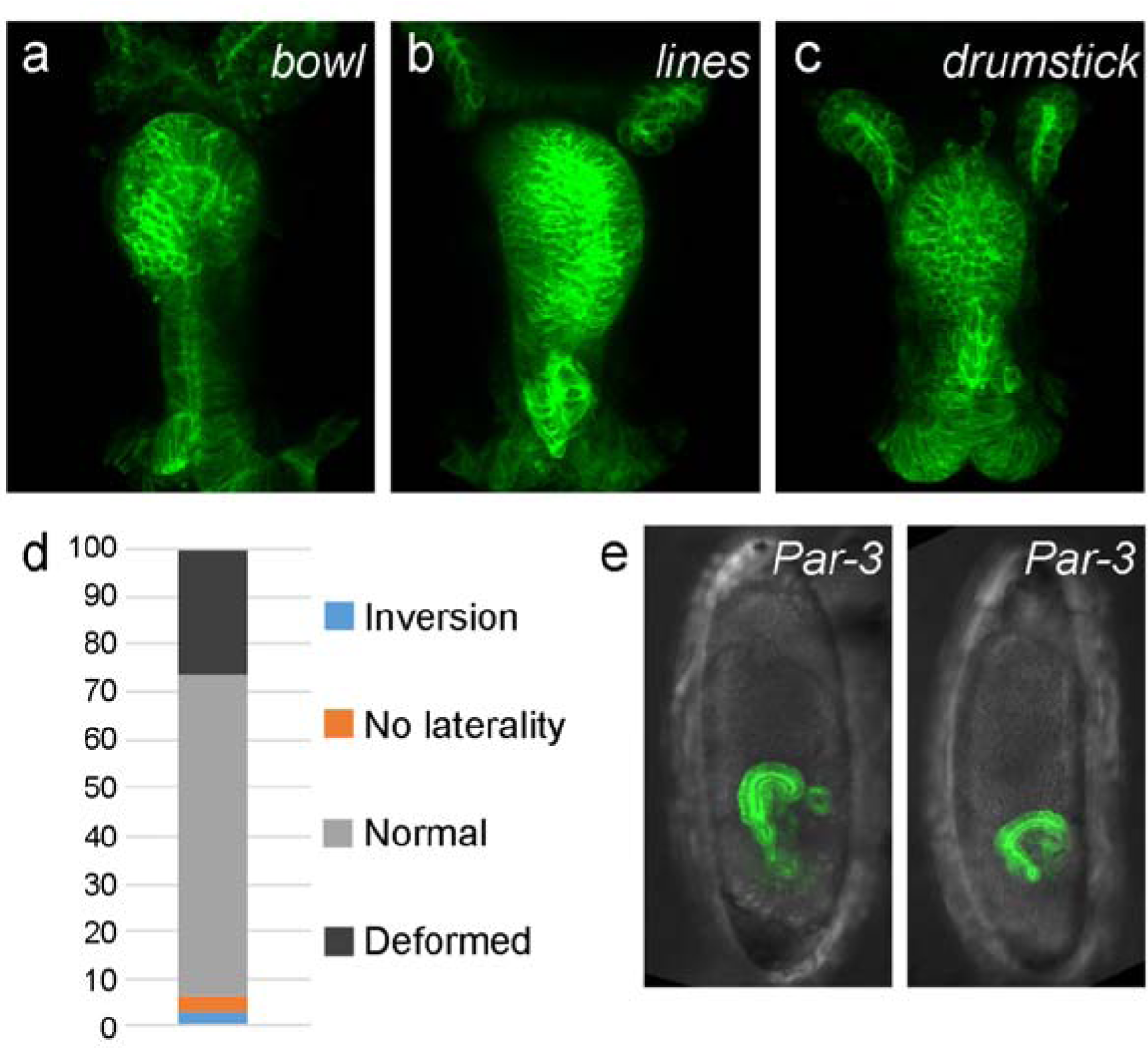
(a-c) The hindgut in bowl (a), lines (b) and drumstick (c) mutants visualized by byn-gal4-driven GFP. All the mutants showed deformation of the hindgut. (d) Frequency of hindgut-shape phenotypes in the *Par-3* mutant embryos. (e) *Par-3* mutant embryos showing delayed germ-band retraction and relatively normal rotation of the hindgut. For all images, anterior is up.

**Figure S3:**
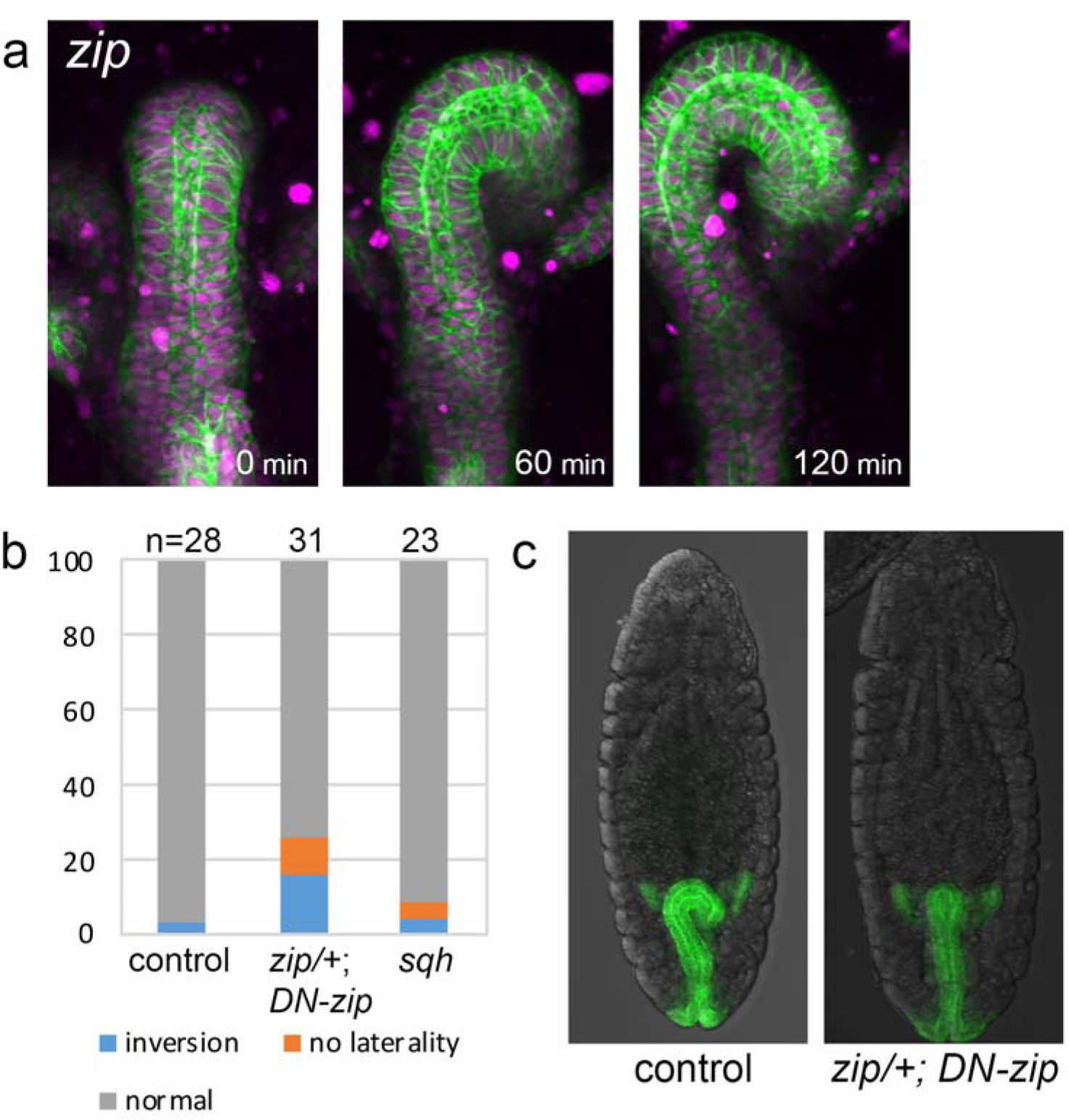
(a) Still shots of the hindgut rotation in *zip* mutant visualized as Fig. 1a. (b) Frequency of hindgut-shape phenotypes in *sqh* mutant and with *zip* disruption. Numbers on the top indicate numbers of examined embryos. (c) The hindgut in wild type (left) and with *zip* disruption (right). Zip-perturbed embryo occasionally showed no rotation of the hindgut. For all images, anterior is up. Elapsed time from the start of the movie is shown at lower right.

